# Ischemic Preconditioning and Left Ventricular Dysfunction: A Novel Mechanism & Model for Pulseless Electrical Activity

**DOI:** 10.1101/468710

**Authors:** Kaustubha D Patil, Richard S Tunin, Sarah J Fink, Susumu Tao, Henry R Halperin

## Abstract

**Background:** Pulseless electrical activity (PEA) is a very common rhythm in cardiac arrest and survival is ≈ 5%. Population data suggest coronary ischemia significantly contributes to PEA, but the mechanism is unknown.

**Objectives:** We hypothesize ischemic preconditioning (IPC) in the setting of left ventricular (LV) dysfunction can convert ischemia-induced ventricular fibrillation (VF) into ischemia-induced PEA.

**Methods:** Using percutaneous coronary interventions in anesthetized swine, we studied the effect of IPC prior to ischemia on arrhythmic burden in normal animals and in animals with LV dysfunction. IPC protocol: four cycles of three minutes of coronary occlusion followed by seven minutes of reperfusion. Chronic LV dysfunction protocol: two serial infarcts in two coronary territories, separated by one week of recovery.

**Results:** In normal animals, IPC prior to ischemia significantly reduced VF incidence (2/8 IPC vs. 7/8 control). In IPC animals with VF, the time to VF was significantly delayed (37.2 ± 7.3 min vs. 20.7 ± 4.9 min, p<0.005). In chronic LV dysfunction animals (EF 15% ± 5%), ischemia caused PEA in all animals (18/18). In non-preconditioned animals, VF followed PEA in all cases (12/12). In preconditioned animals, PEA sustained without VF in 2/6 animals. In 4/6 animals, PEA was prolonged and time to VF was significantly delayed compared to non-preconditioned animals (33.7 ± 7.8 min vs. 12.2 ± 5.0 min, p<0.0001).

**Conclusion:** IPC delays/prevents VF. IPC with LV dysfunction prior to ischemia produces prolonged PEA. IPC with LV dysfunction prior to ischemia is likely an important mechanism for human PEA arrest. This is the first animal model of ischemic pulseless electrical activity.

## Introduction

There are over 500,000 victims of cardiac arrest each year in the United States (300,000 out-of-hospital; 200,000 in-hospital).^1^ In the majority of arrests, the initial rhythm is not ventricular fibrillation (VF) or ventricular tachycardia (VT), but is pulseless electrical activity (PEA) or asystole.^2-5^ The incidence of VT/VF has decreased over the last several decades,^6-10^ while the incidence of PEA/asystole has increased.^1, 8-11^

Data from the National Registry of CPR, reporting on 51,919 in-hospital arrests, found only 24% had an initial rhythm of VT/VF.^5^ The initial rhythm was PEA in 37% and asystole in 39%. Since most patients were monitored and response times were short, the high incidence of PEA/asystole cannot be ascribed to patients being late in cardiac arrest. Similar trends have been seen for out-of-hospital arrests. Data from the Resuscitation Outcomes Consortium, reporting on 12,930 out-of-hospital arrests, found that only 26% had an initial rhythm of VT/VF.^6^ Data from the CARES registry documented similar findings in 25,116 out-of-hospital arrests.^12^ Thus, the majority of cardiac arrests today are due to PEA/asystole. The reasons why there are more PEA/asystolic arrests remain unknown.

The survival rate for VT/VF arrest averages 20%.^1^ The survival rate for PEA/asystolic arrest is substantially lower, averaging 5%.^1, 5^ There is a critical need for improved resuscitation strategies, since each 1% increase in survival rate would result in approximately 5000 additional survivors. This critical need is most apparent in PEA arrest, as little is known about its pathophysiology or its optimal treatment, especially when compared to VT/VF arrest. Understanding PEA has been limited by the lack of a clinically relevant laboratory model.

A substantial number of PEA patients have chronic coronary disease, suggesting that coronary ischemia may be a significant contributing factor.^13^ One study of cardiac arrest found that one-third of patients with resuscitated PEA underwent intervention for acute coronary occlusion.^14^ A major question that remains is why ischemia induces VT/VF arrest in some cases, while inducing PEA arrest in others. In a small percentage of patients, PEA is due to a recognizable and potentially reversible factor (i.e. pulmonary embolus), but specific treatments for these factors have had little effect on survival.^8, 15^ In the vast majority of patients with PEA arrest, the mechanism of the arrest is not well understood.

The effect of ischemic preconditioning (IPC) in the context of ischemia/reperfusion injury has been well studied. ^16-21^ IPC entails exposing myocardial tissue to several short ischemic episodes, followed by recovery, prior to prolonged ischemia. IPC in animal studies significantly mitigates the toxicity of ischemia/reperfusion.^17,20,22,23^ One of these toxic effects is VT/VF, and remarkably, animals preconditioned with short bouts of ischemia prior to prolonged ischemia/reperfusion suffer less VT/VF than non-preconditioned animals.^24-26^

We hypothesized that chronic ischemia induces myocardial conditioning, which prevents or delays the occurrence of VF. We aimed to test whether IPC shifts the effect of a large ischemic burden to cause less VF. We also hypothesized that a large proportion of PEA arrests are due to ventricular muscle failure from acute ischemia in a substrate with chronic left ventricular (LV) dysfunction. We aimed to test whether PEA can result from profound ischemia in the setting of chronic LV dysfunction. We further hypothesized that chronic ischemia in the setting of chronic LV dysfunction shifts the effect of a large ischemic burden from VF to sustained PEA arrest. We aimed to test whether IPC in the setting of chronic LV dysfunction can convert an otherwise ischemia-induced VF arrest into an ischemia-induced PEA arrest.

## Methods

### Study Design

This was a controlled laboratory experiment performed using swine. All animals were treated in accordance with institutional Johns Hopkins Animal Care and Use Committee (Protocol Number: SW16M78) guidelines and in strict compliance with the Animal Welfare Act regulations and Public Health Service Policy. Aortic pressure and electrocardiographic (ECG) monitoring was continuous for all protocols. All hemodynamic tracings included show blood pressure (y-axis) versus time (x-axis). Some included tracings also include a continuous ECG rhythm (y-axis) versus time (x-axis). PEA was defined as mean arterial blood pressure (MAP) below 40 mmHg, if organized ECG complexes were present. Study endpoints included incidence of VF, incidence of PEA, time to VF, and time to PEA.

#### Animal preparation

Pigs (35 ± 5 kg, female, American Yorkshire breed) were tranquilized with sedazine 18 μg/Kg, ketamine 0.9 mg/Kg *IM*, and telazol 0.23 mg/Kg *IM*, anesthetized with pentothal 15 mg/Kg *IV*, intubated and mechanically ventilated (Narkomed 2A, Drager) with 100% O_2_ and 0.5 - 1.5% isoflurane. Percutaneous femoral venous and arterial access was established. An arterial sheath was placed in the femoral artery for systemic blood pressure measurements. Coronary guide catheters (6 Fr) were placed in desired coronary arteries. Angioplasty catheters (3×8mm) were positioned under fluoroscopy in desired coronary arteries.

In some animals, we assayed for coronary reactive hyperemia via Doppler coronary flow wire (Volcano Therapeutics, 0.014” diameter). Transthoracic echocardiograms were performed to assess wall motion, ejection fraction (EF), and stroke volume. All collected hemodynamic and arrhythmia data is shown in Fig 1. Advanced cardiac life support (ACLS) was performed at onset of VF for all animals.

**Fig 1:**
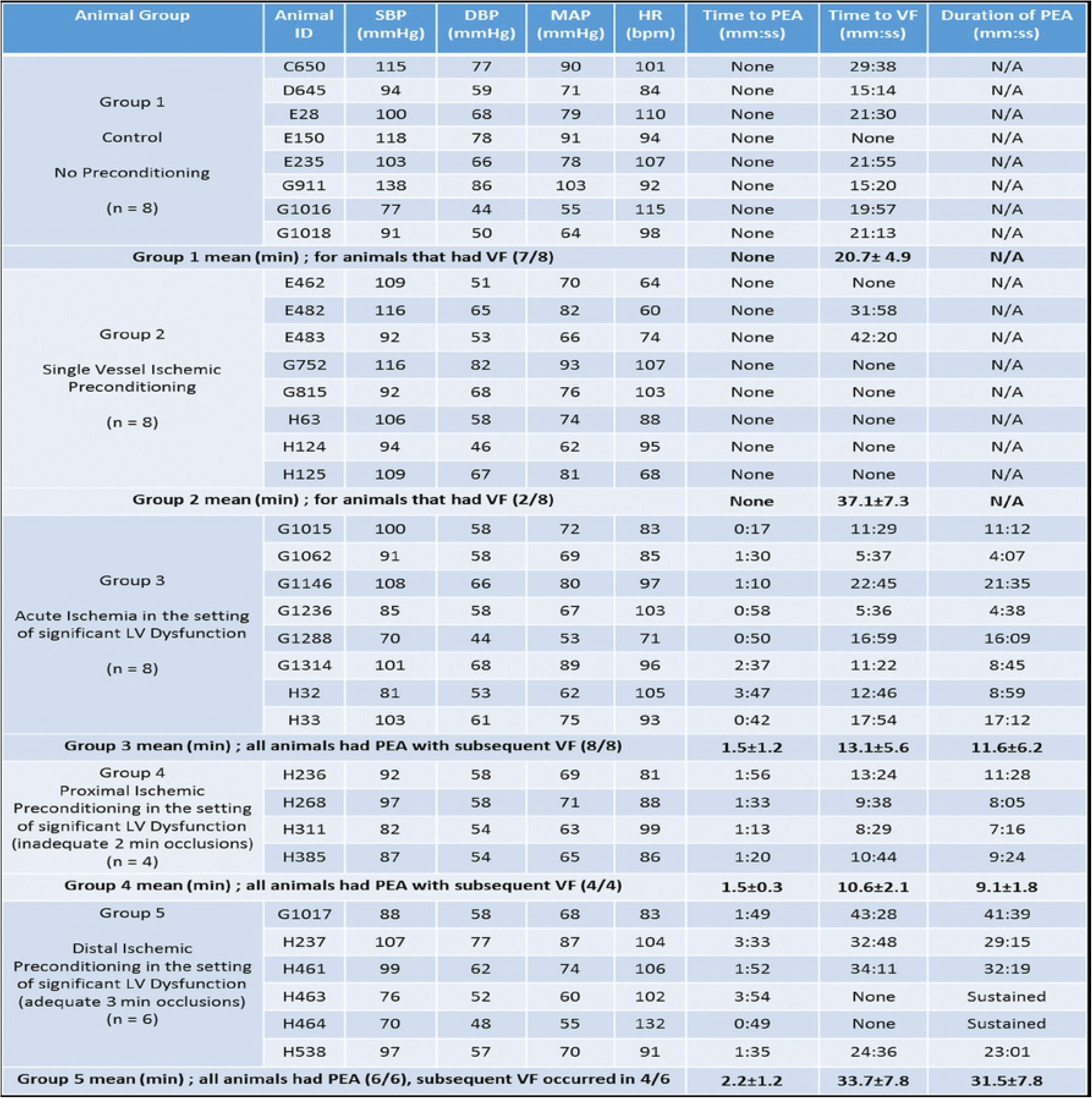
Animal Data: Baseline hemodynamics (in multiple procedures groups, the baseline data listed are prior to the final experiment), Time to PEA, Time to VF, and Duration of PEA (when applicable). Abbreviations: SBP (systolic blood pressure), DBP (diastolic blood pressure), MAP (mean arterial pressure), HR (heart rate)

#### Animal Groups/Protocols (Fig 2)

**Fig 2:**
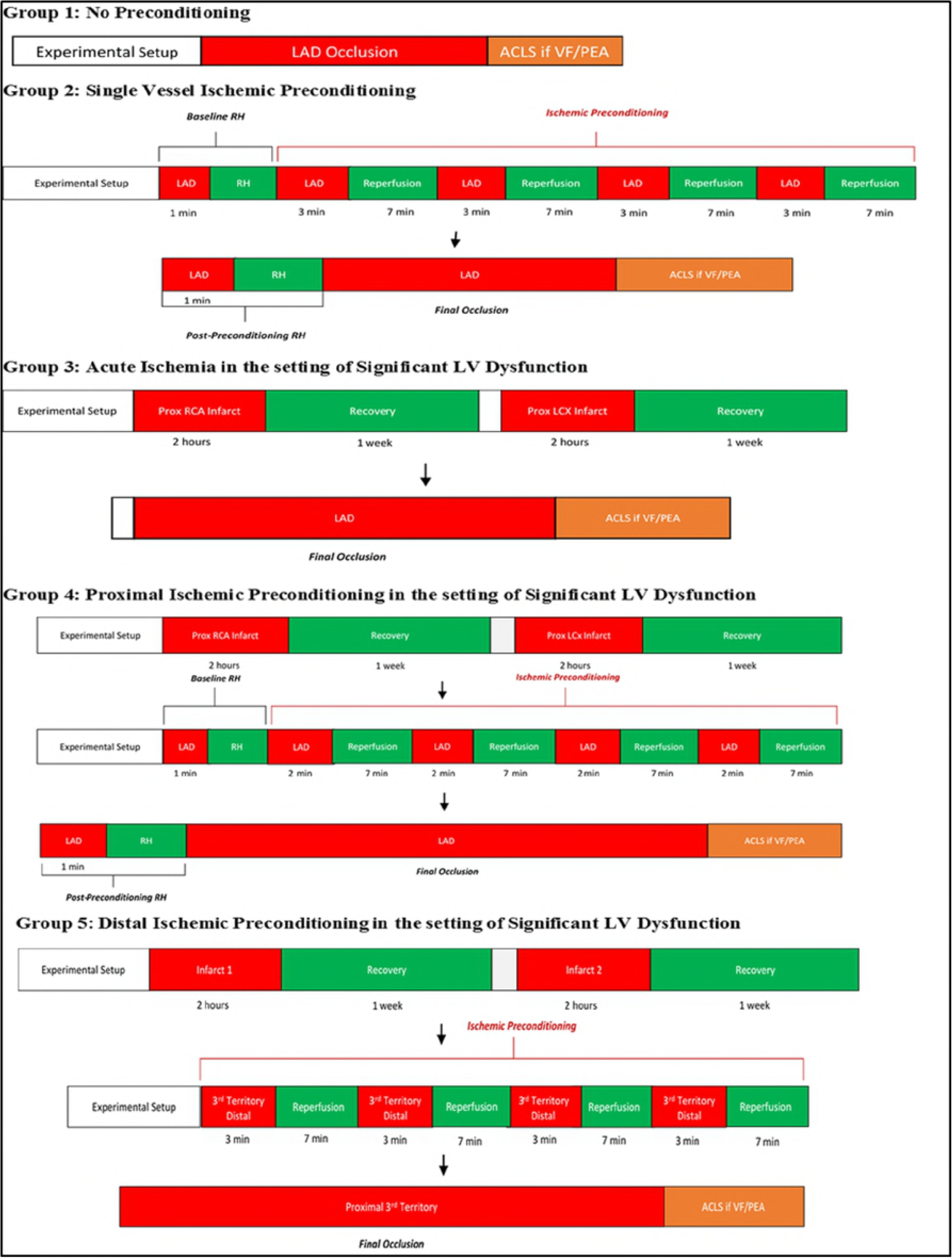
Animal Group Protocols; abbreviations: LAD: left anterior descending, RCA: right coronary artery, LCX: left circumflex artery, ACLS: advanced cardiac life support, RH: reactive hyperemia,VF: ventricular fibrillation, PEA: pulseless electrical activity

##### Acute Ischemia

1. <u>No Preconditioning (n=8)</u>: After the initial preparation, a single mid-left anterior descending (LAD) occlusion was performed for 60 minutes or until VF onset.
2. <u>Single Vessel Ischemic Preconditioning (n=8)</u>: After the initial preparation, four cycles of 3 minutes of occlusion/7 minutes of reperfusion were performed in the LAD territory. After this series, a final mid-LAD occlusion was performed for 60 minutes or until VF onset.

##### Left Ventricular Dysfunction prior to Acute Ischemia

3. <u>Acute Ischemia in the setting of Significant LV Dysfunction (n=8)</u>: Two-hour infarcts were created in both the proximal right coronary artery (RCA) territory and proximal left circumflex (LCX) territory, in that order, by occluding each respective artery for two hours. Infarcts were separated by one week. One week after the LCX infarct, the proximal LAD was occluded for 60 minutes or until VF onset.
4. <u>Proximal Ischemic Preconditioning in the setting of Significant LV Dysfunction (n=4)</u>: Two-hour infarcts were created in both the proximal RCA territory and proximal LCX territory, in that order. Infarcts were separated by one week. One week after the LCX infarct, four cycles of 2 minute occlusion (duration limited by hypotension)/7 minutes of reperfusion were performed in the proximal LAD territory. After this series, a final proximal LAD occlusion was performed for 60 minutes or until VF onset.
5. <u>Distal Ischemic Preconditioning in the setting of Significant LV Dysfunction (n=6):</u> Two-hour infarcts were created choosing either the proximal RCA or mid LAD first, followed by a second infarct in an alternate territory (either proximal LCX or proximal RCA, respectively). Infarcts were separated by one week. One week after the second infarct, four cycles of 3 minutes of occlusion/7 minutes of reperfusion were performed in the distal part of the vessel to the un-infarcted territory. After this series, a final occlusion was performed in the proximal part of the same vessel for 60 minutes or until VF onset.

#### Development of the PEA Model

Before choosing the protocol described in animal group 3, we attempted various sequences of both ischemia and infarct to assess their tendency to lead to VF versus severe hypotension/PEA. This included looking at hemodynamic responses to proximal LAD ischemia only, mid-LAD infarct followed by proximal RCA ischemia only, and proximal RCA infarct followed by proximal LCX ischemia only.

#### Statistical Analysis

Assuming that few, if any, preconditioned animals develop VF, at least four animals/group are needed to show a statistically significant difference compared to control. All mean values are accompanied by standard deviations. All statistical analyses were two-sided and done with alpha = 0.05. Comparisons of means between two groups were performed using unpaired t-test. Survival comparisons between groups were performed using the log-rank test.

## Results

### Acute ischemia in the setting of ischemic preconditioning delays or prevents ventricular fibrillation

#### Animal Group 1: No Preconditioning (n=8)

In this group, 7/8 animals suffered VF cardiac arrest with mean time to VF of 20.7 ± 4.9 min after onset of mid-LAD occlusion. There was no significant hypotension prior to VF in any animals (Fig 3A, 3B).

**Figure 3:**
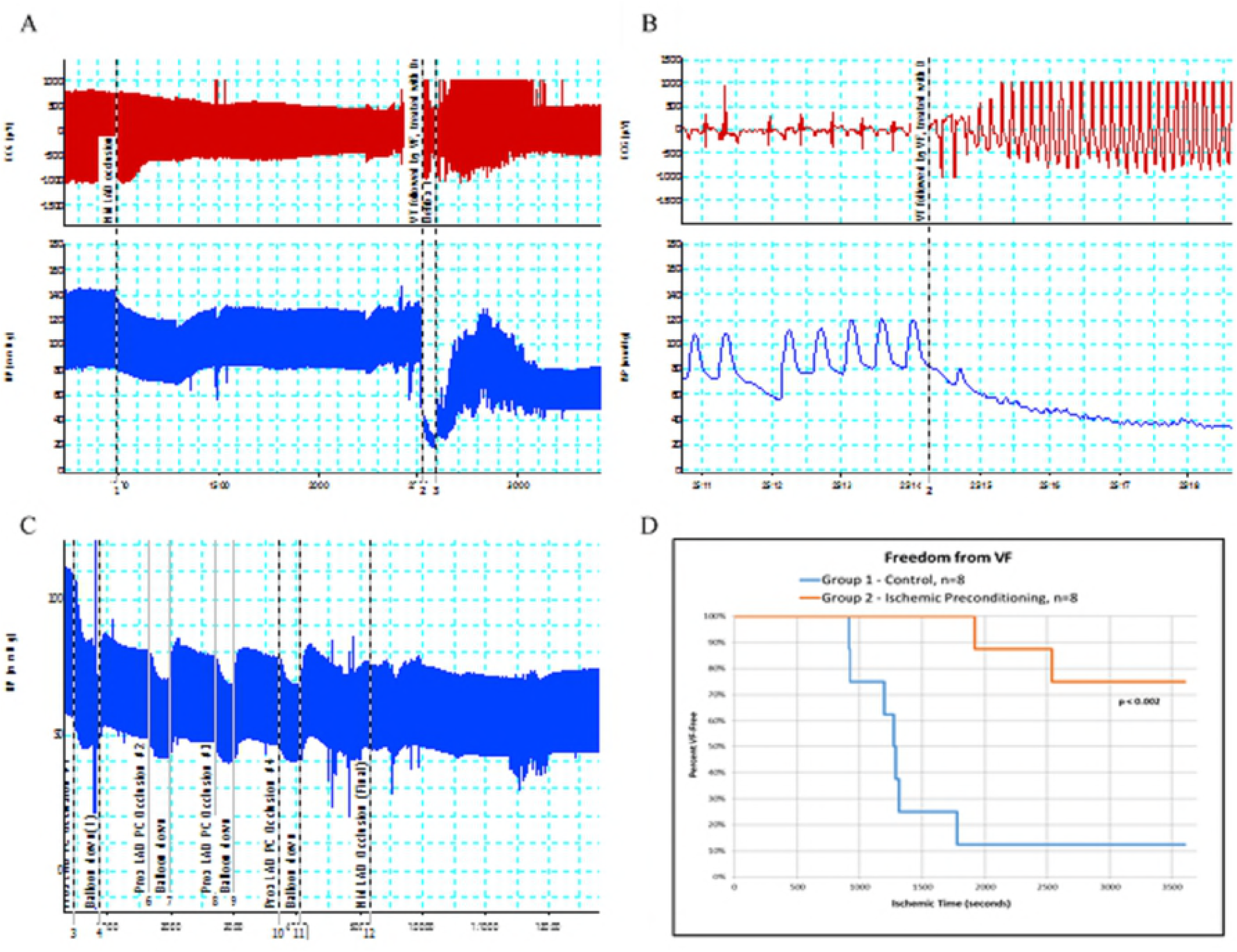
Animals with Acute Ischemia; **A)** Animal Group 1, hemodynamic tracing during mid-LAD occlusion, **B)** Animal Group 1, onset of VF, **C)** Animal Group 2, hemodynamic tracing showing IPC protocol and prolonged final occlusion without VF, **D)** Freedom from VF during ischemia: IPC delays/prevents VF (p < 0.002, log-rank test)

#### Animal Group 2: Single Vessel Ischemic Preconditioning (n=8)

In this group, we tested whether IPC leads to less ischemia-induced VF and thus changes the arrhythmic burden of acute ischemia of a large territory of the heart.

With the IPC protocol, 6/8 animals had no VF during 60 minutes of ischemia. The remaining 2/8 animals had significantly delayed time to VF compared to the 7 animals in group 1 that had VF (37.1 ± 7.3 min vs. 20.7 ± 4.9 min, p < 0.001, t-test). As with group 1 animals, there was no significant hypotension in group 2 animals prior to VF (Fig 3C). Fig 3D compares freedom from VF during ischemia between groups 1 and 2. IPC significantly delayed/prevented VF compared to non-preconditioned animals (p < 0.002, log-rank test).

Although VT/VF suppression is a known effect of IPC (in other species), and could be a marker for successful myocardial preconditioning, we performed an additional assay to confirm successful preconditioning. In prior animal studies (goats and rats), IPC alters the coronary reactive hyperemia response after coronary occlusion.^27-28^ Coronary reactive hyperemia is the transient increase in coronary blood flow upon reperfusion after occlusion of a coronary artery. IPC causes a decrease in time to peak coronary flow upon reperfusion and a decrease in total hyperemic flow.^27^

We used a coronary Doppler flow wire to quantify coronary reactive hyperemia at baseline and after IPC in 10 animals that underwent IPC protocols. A representative coronary flow tracing from one of these animals is shown (Fig 4A). Using area under the curve (AUC) as a surrogate for total hyperemic flow, IPC caused a mean percent decrease of 50% ± 16% in AUC. IPC also caused a mean percent decrease in time to peak coronary flow velocity of 46% ± 14% (Fig 4B). The magnitude of these changes is comparable to the reactive hyperemia response seen in other IPC studies.^27-28^

**Fig 4:**
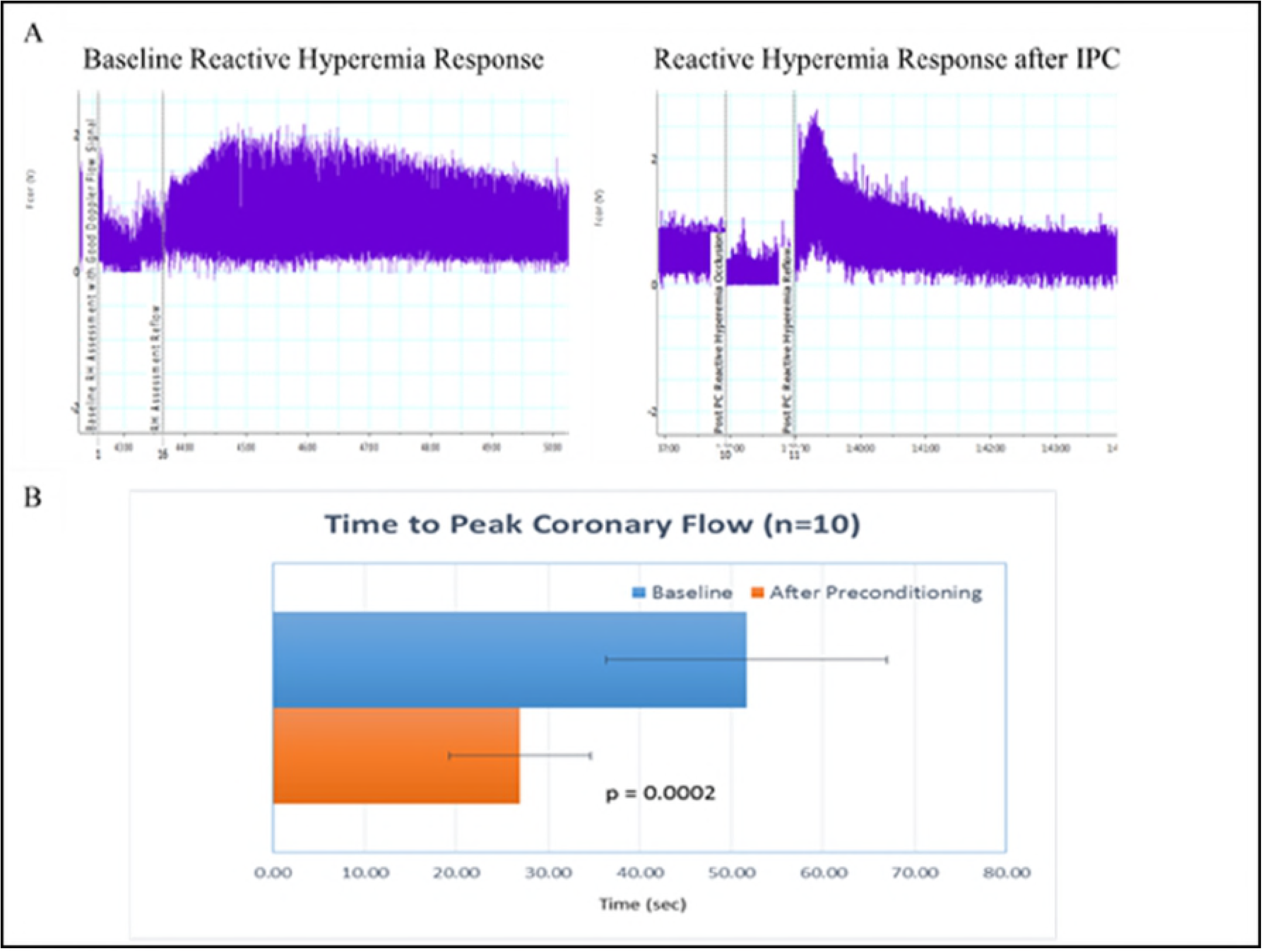
Coronary Reactive Hyperemia; **A)** Coronary flow vs. time during reactive hyperemia at baseline and after IPC. The first vertical line is vessel occlusion and the second is vessel reperfusion. **B)** Time to peak coronary flow velocity during reactive hyperemia at baseline (52±15 sec) and after IPC (27±8 sec), (p <0.001, t-test)

### Developing a model of ischemic PEA arrest

To develop an ischemic model of PEA, we realized that we needed acute ischemia to cause significant hypotension. We postulated that substantial LV dysfunction is necessary prior to acute ischemia for PEA to develop. To this end, we tested multiple protocols of regional coronary ischemia/infarction to assess each protocol’s hemodynamic response to acute ischemia.

#### Proximal LAD Infarct

We first assessed the hemodynamic response to proximal LAD infarct. Although a prolonged proximal LAD occlusion was tolerated, upon reperfusion, animals would invariably suffer VF arrest and resuscitation was rarely successful. Representative hemodynamic tracings are shown (Fig 5A and 5B).

**Fig 5:**
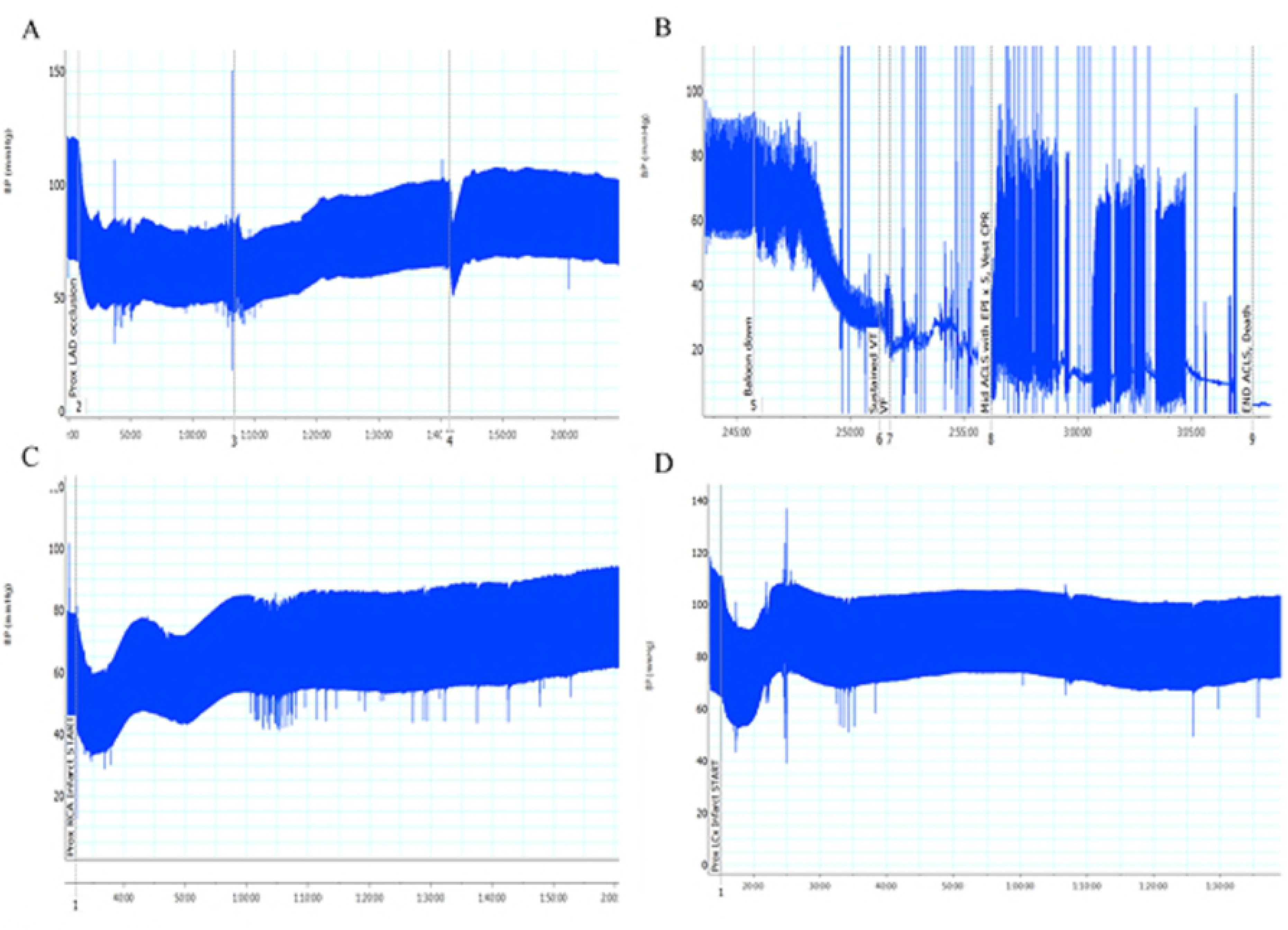
Development of the PEA Model; **A)** Hemodynamic tracing during proximal LAD ischemia, **B)** Hemodynamic tracing showing reperfusion after 2 hours of proximal LAD occlusion. Reperfusion complicated by refractory VF, unable to resuscitate despite 20-25 minutes of ACLS. **C)** Hemodynamic tracing of acute proximal RCA ischemia with prior mid LAD infarction, **D)** Hemodynamic tracing of acute proximal LCX ischemia with prior proximal RCA infarction

#### Proximal Ischemia with Single Prior Infarct

We then looked at the hemodynamic response to acute proximal right coronary artery (RCA) ischemia in the setting of prior mid LAD infarction. These animals tolerated acute RCA ischemia without significant prolonged hypotension/PEA (Fig 5C). Similarly, we looked at the hemodynamic response to acute proximal left circumflex (LCX) artery ischemia in the setting of prior proximal RCA infarction; again, animals tolerated this protocol without significant prolonged hypotension/PEA (Fig 5D).

#### Proximal Ischemia with Multiple Prior Infarcts

We then attempted protocols with multiple prior infarcts. Ultimately, the protocol described below for group 3 animals reliably reproduced hypotension/PEA. This protocol was also the safest for animals to tolerate the multiple infarctions and survive until the final experiment.

### Acute ischemia in the setting of LV dysfunction produces PEA arrest

#### Animal Group 3: Acute Ischemia in the setting of Significant LV Dysfunction (n=8)

In this group, we studied the hemodynamic response to acute ischemia in the setting of prior significant LV dysfunction. Significant LV dysfunction was produced by creating infarcts in both the proximal RCA and proximal LCX territories; infarcts were separated by one week. One week after the LCX infarct, we occluded the proximal LAD. Representative hemodynamic tracings are shown, which include the onset of severe hypotension upon proximal LAD ischemia (Fig 6A), as well as a higher sweep speed showing severe hypotension with organized electrical activity, i.e. PEA (Fig 6B). All 8 animals in this group suffered PEA shortly after LAD occlusion and remained in PEA until subsequent VF. The mean time to VF was 13.1 ± 5.6 min.

**Fig 6:**
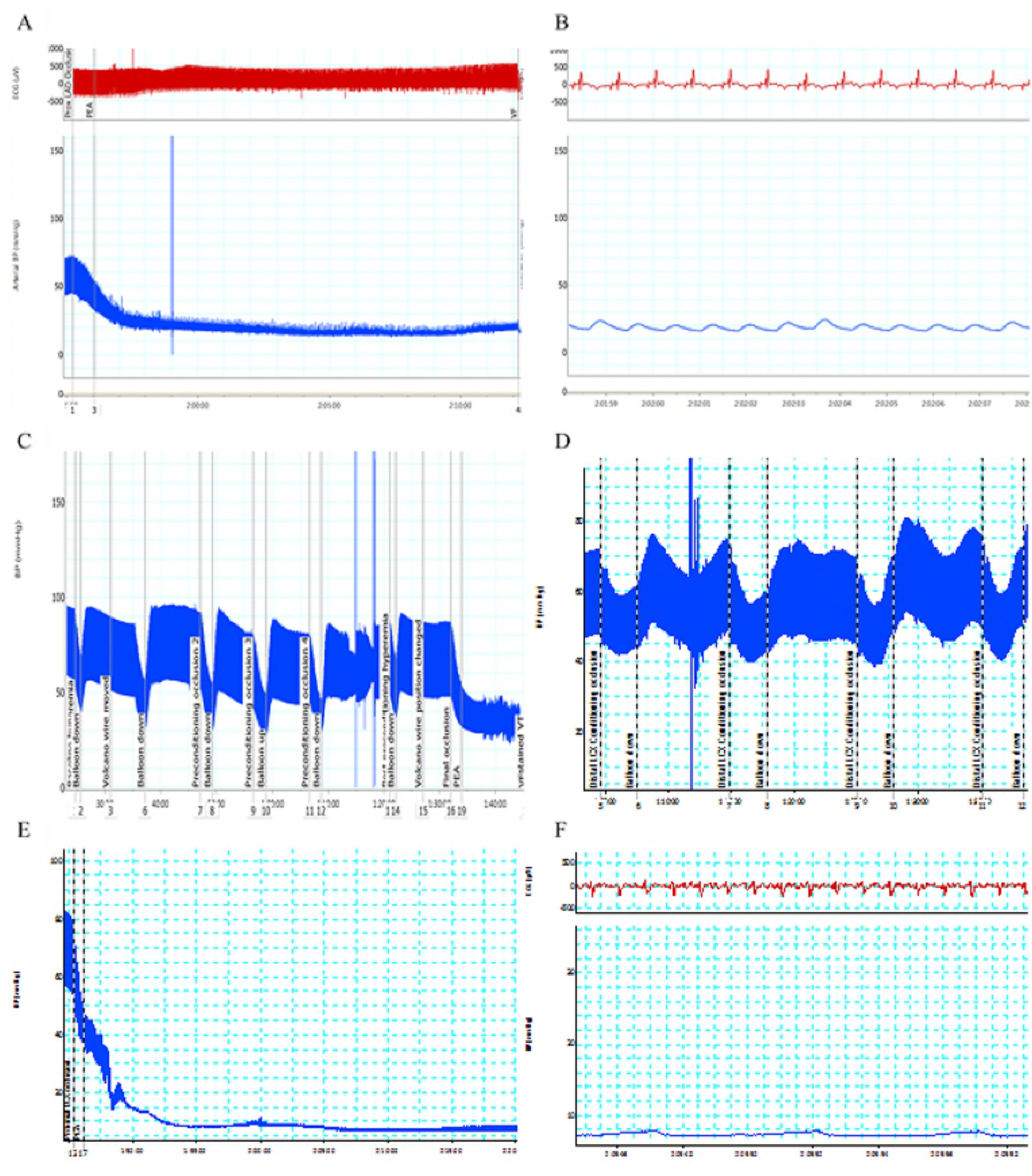
Animals with Left Ventricular Dysfunction prior to Acute Ischemia; **A)** Group 3, hemodynamic response to proximal LAD ischemia in animals with prior LV dysfunction, **B)** Group 3, hemodynamic tracing during successful PEA (severe hypotension with organized ECG complexes), **C)** Group 4, hemodynamic tracing showing IPC (only 2 min occlusions) prior to final proximal LAD occlusion in animals with prior LV dysfunction; note PEA followed by early VF. **D)** Group 5, hemodynamic tracing showing distal LCX IPC (3 min occlusions) followed by reperfusion, **E)** Group 5, acute proximal LCX ischemia in the setting of prior LV dysfunction and distal LCX IPC, **F)** Group 5, *sustained PEA* with severe hypotension and organized ECG (no subsequent VF).

### Acute ischemia in the setting of LV Dysfunction and Ischemic Preconditioning produces sustained PEA arrest

#### Animal Group 4: Proximal Ischemic Preconditioning in the setting of Significant LV Dysfunction (n=4)

Significant LV dysfunction was again created (via proximal RCA and proximal LCX infarctions). Then, we tested whether LAD ischemic preconditioning (remaining viable myocardial territory) prior to acute ischemia can delay or prevent the onset of VF (as in animal group 2). Due to severe hypotension, our proximal LAD IPC occlusions were limited to 2 minutes (instead of 3 minutes as in Group 2 animals). IPC was followed by acute proximal LAD occlusion. We again produced PEA in all animals, and all subsequently had VF arrest at a mean time to VF of 10.6 ± 2.1 min. The mean time to VF was not significantly different from Group 3 (p=0.42, t-test). A representative hemodynamic tracing is shown (Fig 6C).

We suspected that IPC in animal group 4 was unsuccessful in delaying VF due to inadequate preconditioning. IPC was inadequate because the duration of the IPC occlusions in these animals was limited. In these animals, we did not achieve full 3 min IPC occlusions due to severe hypotension during each proximal LAD IPC occlusion.

Furthermore, we wondered whether an alternate sequence of infarctions and final ischemia would alter the propensity for development of PEA. We suspected that the specific territory impacted by the final occlusion was less important than the overall amount of LV dysfunction prior to the final experiment.

#### Animal Group 5: Distal Ischemic Preconditioning in the setting of Significant LV Dysfunction (n=6)

We tested the possibility that short duration IPC (<3 min occlusions) was not successful in delaying VF by adjusting our protocol. We also tested whether the specific final ischemic territory was an important factor in whether or not animals developed PEA during acute ischemia in the context of LV dysfunction. We chose either a mid-LAD infarct or proximal RCA infarct to perform as the first infarct. That was followed one week later by either a proximal RCA infarct (if the first infarct was mid-LAD) or a proximal LCX infarct (if the first infarct was proximal RCA). On the final experiment, however, we performed IPC of the distal remaining territory (either LCX or LAD) to allow four full cycles of 3 min IPC occlusion/7 min reperfusion; MAP was maintained ≥ 45 mmHg during IPC occlusions (Fig 6D). Then, a proximal occlusion was performed in the same un-infarcted territory (for 60 minutes or until VF onset). Representative hemodynamic tracings are shown, which include the onset and duration of PEA upon proximal ischemia (Fig 6E), as well as a higher sweep speed showing organized electrical activity during prolonged PEA (Fig 6F).

In this group, all 6 animals suffered early PEA after final occlusion. In 2/6 animals, there was sustained PEA without subsequent VF for the entire 60 minute occlusion. This compares with 0/12 non-preconditioned animals (groups 3 and 4) that were free from VF during the 60 minute occlusion. In the remaining 4/6 animals in group 5, PEA was prolonged prior to subsequent VF; mean time to VF was significantly delayed compared to non-preconditioned animals (33.7 ± 7.8 min vs. 12.2 ± 5.0 min, p <0.0001, t-test). This protocol successfully produced *sustained* PEA arrest. Distal territory IPC in the setting of significant LV dysfunction significantly delayed or prevented VF during acute ischemia (p<0.001, log-rank test, Fig 7A).

**Fig 7:**
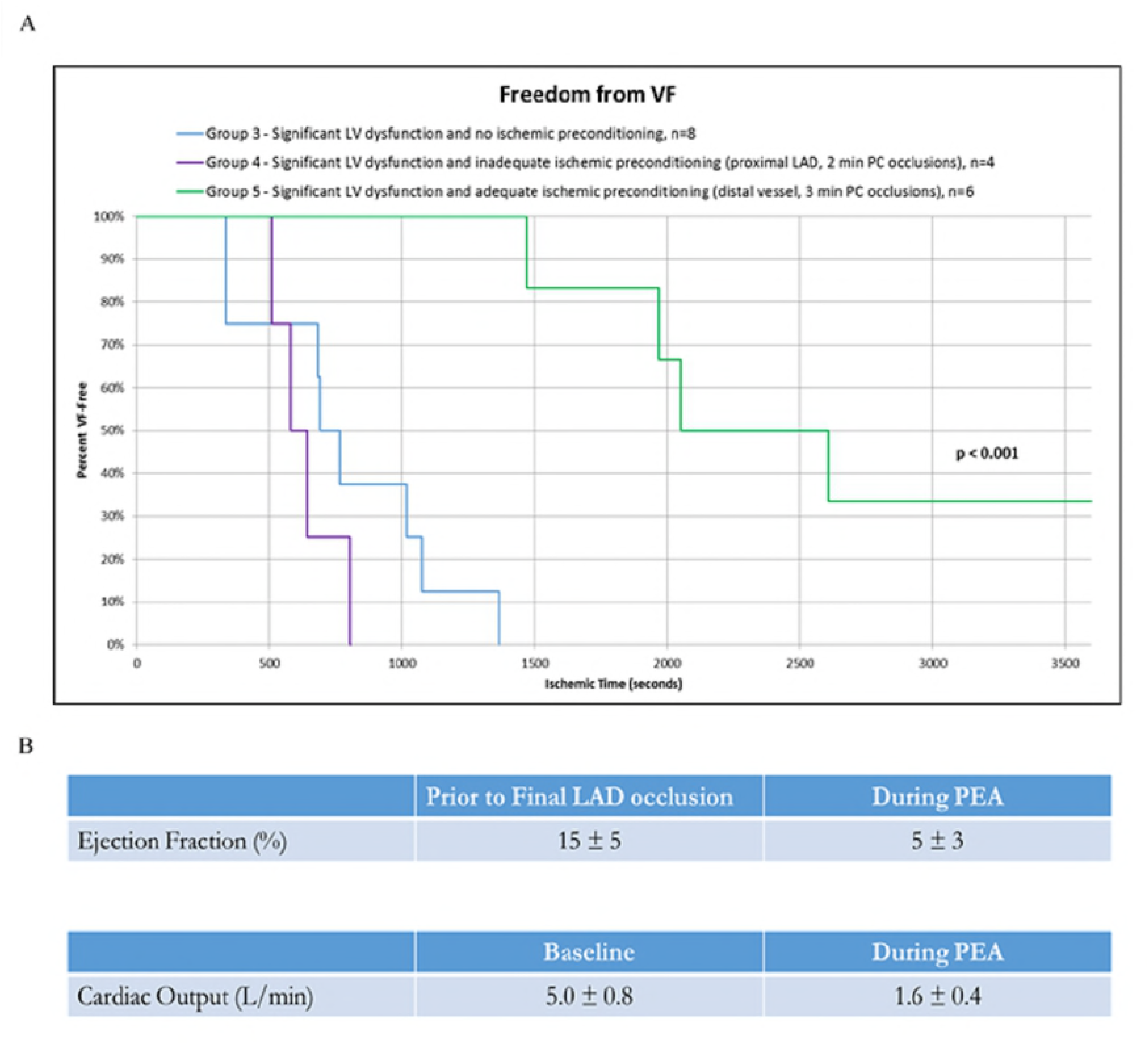
Prolonged PEA; **A)** Freedom from VF during ischemia (in the setting of prior LV dysfunction) comparing animal groups 3-5: IPC (3 min occlusions in a distal territory) successfully delayed/prevented VF and led to *sustained* PEA after final ischemia in the setting of prior LV dysfunction (p < 0.001, log-rank test) **B)** Severe cardiac dysfunction is necessary for PEA (n = 12)

LV dysfunction (via transthoracic echocardiography) was quantified in most animals with PEA arrest (n=12). A normal LV EF at baseline in swine is similar to that in humans, approximately 50-65% (Supplemental Movies S1-2). The amount of cardiac dysfunction necessary for PEA was substantial (Fig 7B). Mean EF prior to the final experiment was 15% ± 5%, consistent with our aim of producing significant LV dysfunction prior to the final ischemic insult (Supplemental Movies S3-4). The mean EF necessary for PEA onset was 5% ± 3% (Supplemental Movies S5-6). Furthermore, the cardiac output (CO) necessary for PEA onset was 1.6 ± 0.4 L/min, approximately a 70% reduction from baseline (CO of 5.0 ± 0.8 L/min).

## Discussion

There are three major new findings in this study: 1) ischemic preconditioning delays or prevents the onset of VT/VF in a swine model of acute coronary ischemia, 2) acute ischemia in the setting of substantial, chronic LV dysfunction reliably produces PEA cardiac arrest, and 3) ischemic preconditioning in the setting of substantial, chronic LV dysfunction reliably produces *sustained* PEA arrest upon acute ischemia.

We demonstrated that IPC delays/prevents the onset of VT/VF in a swine model of acute coronary ischemia (Fig 3D).

Our goal however, was to model clinical PEA. Traditionally, PEA is classified as normotensive PEA, pseudo-PEA, and true PEA.^45^ Normal myocardial contractions in the absence of detectable pulses is “normotensive” PEA. This occurs secondary to a reversible condition, i.e. tension pneumothorax, tamponade, or hypovolemia. Normotensive PEA does not represent a significant subset of patients with clinical PEA. Pseudo-PEA is characterized by weak myocardial contractions that produce detectable aortic pressure only measurable by invasive monitoring or echocardiography.^29-31^ True PEA is the absence of myocardial contractions, typically the final stage of PEA occurring after prolonged exposure to acidosis and hypoxia.^29-31^

Pseudo-PEA represents a substantial portion of PEA arrest^31^ and population data suggest coronary ischemia is a major contributor to PEA.^12^ Thus, we attempted to create an ischemic model of PEA. If ischemia is to cause PEA, it needs to produce weak myocardial contractions unable to achieve an adequate perfusion pressure despite organized cardiac electrical activity.

With mid-LAD ischemia (animal groups 1-2), significant hypotension was not observed prior to VF (despite delaying/preventing VF in group 2). We proposed that significant LV dysfunction prior to acute ischemia may produce the desired hypotension and thus, PEA. This approach to modeling PEA is clinically relevant (patients with PEA often have underlying cardiomyopathy and LV dysfunction).

### Development of the PEA Model

To create LV dysfunction, we tested various ischemia/infarction sequences to quantify the dysfunction required to induce the necessary hypotension. We attempted single vessel proximal ischemia (LAD) as well as single infarcts followed by alternate vessel ischemia. Neither of these protocols produced enough hypotension.

A two-infarct protocol prior to prolonged ischemia in the third coronary territory finally reliably induced severe hypotension/PEA. This is the second major finding in this study: prolonged ischemia in the setting of significant LV dysfunction reliably produces PEA. This is the first animal model of ischemic PEA cardiac arrest. PEA was reliably induced, however it was soon followed by subsequent VF (animal group 3).

In human adult PEA arrest, PEA is often prolonged (> 20 minutes) prior to discontinuation of resuscitation efforts if there is no return of spontaneous circulation. Next, we aimed to model the *prolonged* PEA that is a common clinical scenario. Perhaps prior LV dysfunction and ischemic preconditioning prior to prolonged ischemia could delay/prevent VT/VF and lead to sustained PEA?

Animal groups 4 and 5 investigated this hypothesis and successfully showed that LV dysfunction and ischemic preconditioning prior to acute ischemia reliably produced *sustained* PEA. Our results show that a critical duration of ischemia is necessary for successful IPC (to exert an anti-arrhythmic effect upon prolonged ischemia). Furthermore, we were successful in delaying VF (animal group 5) despite only exposing a distal coronary territory to IPC. IPC was successful via “remote” preconditioning. That is, the entire territory (and likely the entire myocardium) was conditioned despite a distal site of IPC. In animals with prior mid-LAD and proximal RCA infarcts, where only the distal LCX territory underwent IPC, it follows that all of the remaining un-infarcted territory (supplied by the LCX and proximal LAD) was preconditioned given that we achieved sustained PEA without early VF upon acute ischemia. Remote IPC conferring global myocardial protection from ischemia has been reported. ^26, 32-34^

Given the success of remote IPC, we attempted a hyper-acute (single experiment) model of sustained PEA: a proximal RCA infarct was created, then immediate distal LAD IPC, and finally proximal LAD occlusion. Early VF occurred without PEA; non-sustained ventricular ectopy was noted throughout after the RCA infarct. We suspect the underlying pathobiology of acute infarct (followed by reperfusion) led to increased arrhythmogenicity and predisposition to VF. This confirmed our hypothesis that *chronic*, not acute, LV dysfunction is necessary for ischemic PEA.

Severe cardiac dysfunction is necessary for PEA (mean EF ~15% prior to final prolonged ischemia, mean EF ~5% at PEA onset). PEA onset required approximately a 70% reduction in cardiac output from baseline. We suspected that LV dysfunction was necessary to induce PEA; however, this severity of dysfunction was beyond our expectations.

### The Role of Ischemic Preconditioning in Human PEA/Clinical Implications

Our work offers important and novel insights into the pathophysiology and development of human PEA cardiac arrest.

Many PEA arrest patients have chronic coronary disease^12-14^, and we suspect these patients experience numerous ischemic episodes over their lifetime via unstable coronary syndromes, stable angina, and asymptomatic small vessel ischemia. Chronic ischemia may induce myocardial conditioning similar to the effect of IPC in our model. IPC has a substantial anti-arrhythmic effect, and chronic ischemic events (in addition to the antiarrhythmic effects of cardiovascular medications including beta-blockers^35^) may prevent ischemic VF arrest and result in both symptomatic infarcts and so-called “silent” infarcts. Infarcts contribute to progressive LV dysfunction, which likely predisposes patients to PEA arrest with their next ischemic insult.

Even distal territory ischemia promoted global myocardial preconditioning and exerted an impressive anti-arrhythmic effect in our model. Many patients with coronary disease suffer from chronic small vessel ischemia that is asymptomatic and undiscovered. When discovered, revascularization is often contraindicated due to vessel size. This provides the perfect substrate for global ischemic preconditioning and promotes LV dysfunction.

This is the third major new finding of this study: chronic ischemia and chronic LV dysfunction provide the critical substrate and mechanism by which acute ischemia can convert an otherwise ischemia-induced VF arrest into an ischemia-induced PEA arrest (Fig 8).

**Fig 8:**
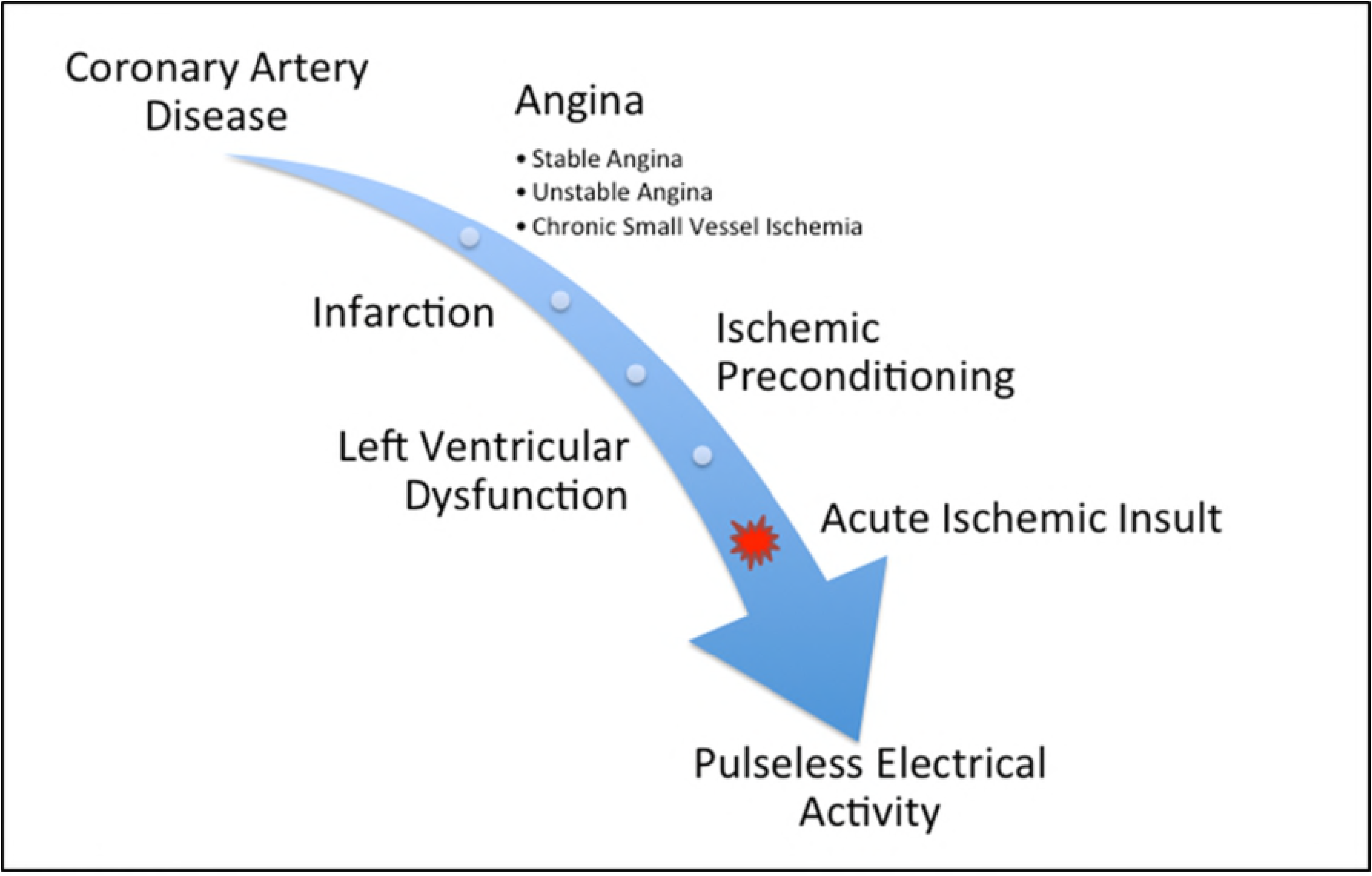
Ischemic Preconditioning and Left Ventricular Dysfunction: a novel mechanism for ischemic pulseless electrical activity. Coronary artery disease and chronic ischemia (via stable angina, unstable angina, and asymptomatic small vessel ischemia) cause ischemic preconditioning, which increases the likelihood of arrhythmia-free survival from infarction. Infarction leads to progressive left ventricular dysfunction and with ongoing ischemic preconditioning, provides the necessary substrate for the development of pulseless electrical activity with the next profound ischemic insult.

It should be no surprise then, that as incident heart failure continues to rise and more people with chronic coronary disease are living longer with more severe ischemic disease (due to improved therapies),^36^ that the proportion of PEA cardiac arrest has also risen concomitantly over the last half century.^4-13^

## Conclusions and Future Directions

Ischemic preconditioning can delay or prevent the onset of ventricular tachyarrhythmias in a swine model of acute coronary ischemia. A profound, acute, ischemic insult on a background of chronic LV dysfunction reliably produces PEA arrest. This is the first animal model of ischemic PEA arrest. Furthermore, ischemic preconditioning prior to acute ischemia in the context of chronic LV dysfunction reliably produces *sustained* PEA arrest, similar to human PEA arrest. This is a novel pathophysiologic mechanism for ischemia-related PEA arrest.

Substantial cardiac dysfunction is necessary for ischemic PEA. Despite the severity of LV dysfunction, animals were doing well (by veterinary metrics, including responsiveness, activity level, and oral intake) prior to the acute ischemic insult that induced PEA. This suggests that PEA may also be reversible acutely. However, given the profound cardiac dysfunction necessary for PEA, reversal of PEA likely requires a resuscitation algorithm that is just as profound and/or aggressive. The currently practiced resuscitation algorithm is likely inadequate to successfully treat PEA cardiac arrest, leading to today’s abysmal survival rates.

We aim to use this ischemic PEA model to study the effects of novel and aggressive techniques in the optimization of systemic/cerebral blood flow during cardiopulmonary resuscitation, because improved blood flow leads to enhanced survival during cardiac arrest.^37-42^

## Limitations

The main limitation of applying our study to human health is that swine are more prone to ventricular arrhythmias in response to coronary ischemia than humans.^43^ Thus, the reproducibility of VF shown in our non-preconditioned animals may not be directly generalizable to human coronary ischemia/VF. However, demonstrating that we can delay or prevent VF in swine despite their susceptibility to arrhythmia is a significant finding.

## Acknowledgements

We thank the Johns Hopkins Animal Care and Use Committee for their support with animal husbandry.

## Supplementary Digital Content Captions

**Movie S1.** Baseline transthoracic echocardiogram – parasternal long axis view

**Movie S2.** Baseline transthoracic echocardiogram – parasternal short axis view

**Movie S3.** Pre-Final Occlusion; Chronic Left Ventricular Dysfunction – parasternal long axis view

**Movie S4.** Pre-Final Occlusion; Chronic Left Ventricular Dysfunction – parasternal short axis view

**Movie S5.** Pulseless Electrical Activity (PEA) during final occlusion – parasternal long axis view

**Movie S6.** Pulseless Electrical Activity (PEA) during final occlusion – parasternal short axis view

## Abbreviations

VT: ventricular tachycardia
VF: ventricular fibrillation
PEA: pulseless electrical activity
IPC: ischemic preconditioning
LV: left ventricle
LAD: left anterior descending
RCA: right coronary artery
LCX: left circumflex
ACLS: advanced cardiac life support
RH: reactive hyperemia
ECG: electrocardiogram
MAP: mean arterial pressure

